# Microglial CX3CR1^I249/M280^ variant limits neurogenesis and remyelination in cuprizone-induced multiple sclerosis model

**DOI:** 10.1101/2021.06.06.447262

**Authors:** Andrew S. Mendiola, Kaira A. Church, Sandra M. Cardona, Difernando Vanegas, Shannon A. Garcia, Wendy Macklin, Sergio A. Lira, Richard M. Ransohoff, Erzsebet Kokovay, Chin-Hsing Annie Lin, Astrid E. Cardona

## Abstract

Microglia have been implicated in multiple sclerosis (MS) pathogenesis. The fractalkine receptor CX3CR1 regulates the activation of pathogenic microglia in models of MS and the human polymorphic *CX3CR1*^*I249/M280*^ (h*CX3CR1*^*I249/M280*^) variant increases MS disease progression. However, the role of h*CX3CR1*^*I249/M280*^ on microglial activation and central nervous system repair and regenerative mechanisms remain unknown. Therefore, using transgenic mice expressing the h*CX3CR1*^*I249/M280*^ variant, we aimed to determine the contribution of defective CX3CR1 signaling to remyelination and neurogenesis in the cuprizone model of focal demyelination. Here, we report that mice expressing h*CX3CR1*^*I249/M280*^ exhibit marked demyelination and microgliosis follow acute cuprizone treatment. Cuprizone-treated CX3CR1-deficient and fractalkine-deficient mice displayed a comparable phenotype. Nanostring gene expression analysis in demyelinated lesions showed that h*CX3CR1*^*I249/M280*^ upregulates genes associated with inflammation, oxidative stress and disease-associated microglia. In addition, gene expression analysis in the subgranular zone (SGZ) of the hippocampus in h*CX3CR1*^*I249/M280*^ mice was associated with a significant downregulation of gene networks linked to neurogenesis following acute demyelination. Confocal microscopy showed that h*CX3CR1*^*I249/M280*^ or loss of CX3CR1 signaling inhibits the generation of progeny from the neurogenic niche, including cells involved in myelin repair. These results provide evidence for the pathogenic capacity of h*CX3CR1*^*I249/M280*^ on microglia dysfunction and therapeutic targeting of CX3CR1 to promote CNS repair in MS.

## Introduction

Chronic activation of microglia is associated with MS disease progression (Absinta et al., 2020), and it is has been suggested that resolution of inflammatory microglial responses following demyelination is critical for CNS repair, including remyelination and neurogenesis (Miron et al., 2013; Zhang et al., 2020). However, the molecular mechanisms coupling pathogenic microglia to impaired remyelination and neurogenesis in CNS inflammatory lesions remain poorly defined. Identifying upstream molecular circuits governing microglia-mediated neuroinflammation may provide novel therapeutic strategies for improving motor deficits and cognition in MS.

Interaction of fractalkine, a neuron-derived chemokine, with CX3CR1 inhibits neurotoxic microglia in models of Parkinson’s disease, amyotrophic lateral sclerosis and MS (Cardona et al., 2006; Garcia et al., 2013). It has been shown that CX3CR1 signaling limits demyelination (Garcia et al., 2013) and aberrant remyelination (Lampron et al., 2015) in MS models. Additional evidence supports that CX3CR1 signaling promotes the maturation of oligodendrocyte precursor cells during development (Voronova et al., 2017). In humans, the variant hCX3CR1^I249/M280^ arises from two single-nucleotide polymorphisms in linkage disequilibrium in the CX3CR1 loci and is identified in >20% of some ethnic groups. This mutant receptor is proposed to function as a hypomorphic and as a dominant-negative allele, such that cells expressing the variant are less responsive to fractalkine (McDermott et al., 2003). Expression of hCX3CR1^I249/M280^ is associated with secondary progressive MS in which patients develop more demyelinated lesions (Stojković et al., 2012) and mice expressing hCX3CR1^I249/M280^ exhibit more severe clinical disease in EAE (Cardona et al., 2018). However, the contribution of hCX3CR1^I249/M280^on the inflammatory programming of microglia and its role in myelin repair and formation of new neurons is not known.

In this study, we characterized the hCX3CR1^I249/M280^ variant in response to demyelination and identified that defective CX3CR1 signaling drive neurotoxic microglia by global activation of inflammatory and prooxidant gene programs. We show that this microglial dysfunction mediated by hCX3CR1^I249/M280^ variant suppresses transcriptional and protein markers of neurogenesis. Data also demonstrate that hCX3CR1^I249/M280^ indirectly impairs remyelination by inhibiting oligodendrocyte differentiation following cuprizone treatment. These findings underscore the neuroprotective effects of fractalkine and examination of the exact mechanism by which fractalkine/CX3CR1 regulates the formation of new neurons and myelin-producing cells may provide novel pathways to enhance CNS tissue repair.

## Results

### hCX3CR1^I249/M280^ variant exacerbates demyelination and microgliosis in cuprizone model

To investigate the contribution of defective CX3CR1 signaling through hCX3CR1^I249/M280^ variant on demyelination, we compared the effects of acute cuprizone-induced demyelination (4 wks treatment) in hCX3CR1^I249/M280^, CX3CR1-KO and CX3CR1-WT mice (Fig. 1). Confocal microscopy of myelin immunostaining against MBP and PLP showed that hCX3CR1^I249/M280^ mice were highly sensitive to cuprizone-induced demyelination with exacerbated lesions in the corpus callosum relative to CX3CR1-KO and CX3CR1-WT mice (Fig. 1A; white arrows). Immunohistochemistry of total myelin confirmed demyelination patterns in all genotypes (Fig. 1A; Blackgold staining). By assessing myelin immunoreactivity throughout the corpus callosum, the extent of demyelination was more evident in the anterior of corpus callosum in hCX3CR1^I249/M280^ mice (38 ± 13 % myelin remaining in corpus callosum) compared to both CX3CR1-WT and –KO mice (66 ± 13 and 54 ± 13 % myelin remaining in corpus callosum, respectively) (Fig. 1B, C).

**Figure 1.**
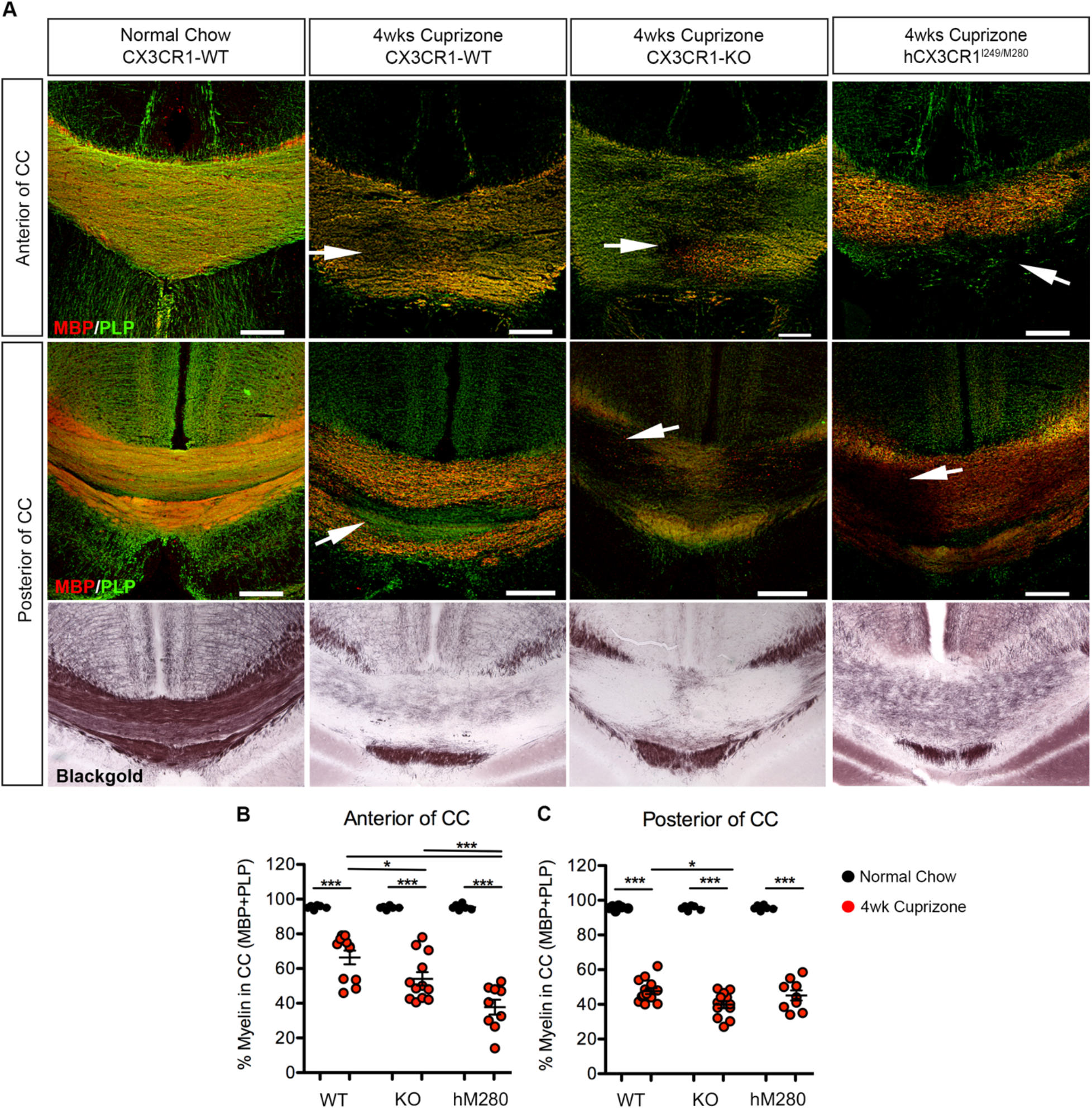
The hCX3CR1^I249/M280^ polymorphism exacerbates cuprizone-induced demyelination. **A**, Representative confocal images of brain sections from normal-chow fed and 4 wk cuprizone fed mice, immmunostained for MBP (red) and PLP (green). White arrows denote demyelinated lesions in the corpus callosum. Immunohistochemistry of the corpus callosum stained with blackgold total myelin stain is shown at the bottom panel. **B, C**. Image quantification of MBP+PLP myelin staining in the (**B**) anterior and (**C**) posterior of the corpus callosum from 4wks cuprizone (red-filled symbols) and normal-chow fed (black-filled symbols) mice. WT, CX3CR1-WT; KO, CX3CR1-KO; hM280, hCX3CR1^I249/M280^. Mean ± SEM, *n* = 6 to 12 mice per group; each dot represents an individual mouse. *P<0.05, ***<0.001 using one-way ANOVA followed by Tukey’s posttest. Scale bars: 200 μm

Confocal microscopy of the microglia/macrophage marker IBA1 revealed a significant increase in microgliosis throughout the corpus callosum of cuprizone-treated hCX3CR1^I249/M280^ and CX3CR1-KO mice relative to CX3CR1-WT mice (Fig. 2A, B). However, microglia densities were more prominent in cuprizone-treated hCX3CR1^I249/M280^ mice as evident by a 1.8-fold increase in IBA1 immunoreactivity compared to cuprizone-treated CX3CR1-WT mice. Because microglia recruitment and phagocytosis of myelin debris is essential for CNS repair (Kotter et al., 2006), we, therefore, assessed well-known microglia activation markers associated with efficient myelin clearance in the cuprizone model (Olah et al., 2012; Cantoni et al., 2015). qPCR analysis showed that acute cuprizone treatment induced a significant increase in *Cxcl10, Cd68*, and *Trem2* gene expression in the corpus callosum of hCX3CR1^I249/M280^ mice compared to both CX3CR1-WT and –KO mice (Fig. 2C). Together, these findings show aberrant microglial responses and augmented CNS lesion formation in hCX3CR1I249/M280 mice. These data are consistent with a previous report showing that hCX3CR1^I249/M280^ promotes EAE clinical severity with pathogenic activation of microglia (Cardona et al., 2018). Areas of active demyelination colocalized with microglial clusters (Figs. 1A, and 2A).

**Figure 2.**
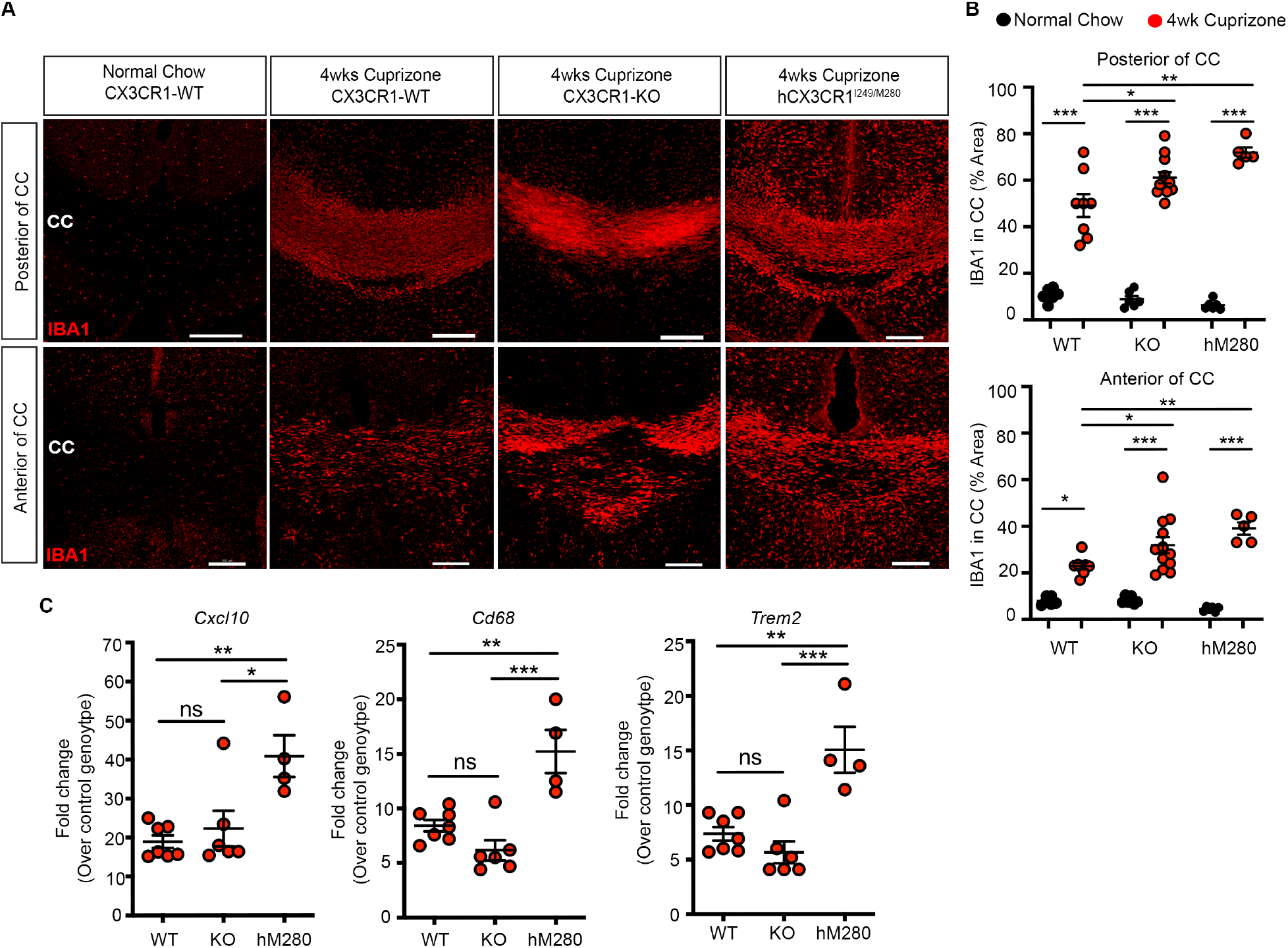
hCX3CR1^I249/M280^ increases microgliosis and activation during cuprizone-induced demyelination. **A**, Representative confocal images of brain sections from normal-chow fed and 4 wk cuprizone fed mice, immmunostained for the microglial marker IBA1 (red), in the posterior and anterior of the corpus callosum of 4wk cuprizone-treated hCX3CR1^I249/M280^, CX3CR1-KO (KO) and CX3CR1-WT mice. Scale bars: 200 μm. **B**, Image quantification of IBA1-positive microglia in the posterior and anterior of the corpus callosum from mice fed normal chow (black-filled symbols) or 4wks cuprizone (red-filled symbols). Mean ± SEM, *n* = 6 to 12 mice per group, each dot represents an individual mouse. **C**, Quantitative PCR analysis of *Cxcl10, Cd68*, and *Trem2* gene expression in corpus callosum isolated from mice treated with or without cuprizone. Data are expressed as mean fold change over respective normal-chow fed genotype ± SEM, *n* = 4 to 7 mice per group; each dot represents an individual mouse. *P<0.05, **P<0.01, ***<0.001 using one-way ANOVA followed by Tukey’s posttest. ns= not significant.

### Loss of fractalkine mimics hCX3CR1^I249/M280^ phenotype in cuprizone model

To test if loss of fractalkine, the only known CX3CR1 ligand, was sufficient to mimic the demyelinating phenotype in cuprizone-treated CX3CR1-KO and hCX3CR1^I249/M280^ mice, we treated FKN-KO (fractalkine deficient) mice (Cook et al., 2001) with cuprizone. FKN-KO mice treated with cuprizone showed a significant reduction in myelin in the corpus callosum compared to normal chow controls (Fig. 3A-C). The level of demyelination in FKN-KO mice was comparable to that observed in cuprizone-treated CX3CR1-KO and hCX3CR1^I249/M280^ mice (Fig. 1). FKN-KO mice showed robust microgliosis and activation in both the anterior and posterior regions of the corpus callosum following cuprizone treatment (Fig. 3D-F). In cuprizone-treated FKN-KO mice, confocal microscopy of IBA1+ microglia showed activated cells with typical amoeboid morphology, and truncated cellular processes (Ransohoff, 2016), colocalizing with increased expression of the phagocytic marker CD68. (Fig. 3D-F). As expected, FKN-KO mice fed normal chow displayed microglia with a ramified morphology and basal levels of CD68 expression (Fig 3D, E, lower panels; yellow arrows). Image quantification showed cuprizone led to a significant increase in both IBA1 and CD68 immunoreactivity in the corpus callosum (Fig. 3F), consistent with the microgliosis observed in both CPZ-treated CX3CR1-KO and hCX3CR1^I249/M280^ mice (Fig. 2). These results suggest that fractalkine-CX3CR1 signaling limits microglia activation and demyelination following acute cuprizone.

**Figure 3.**
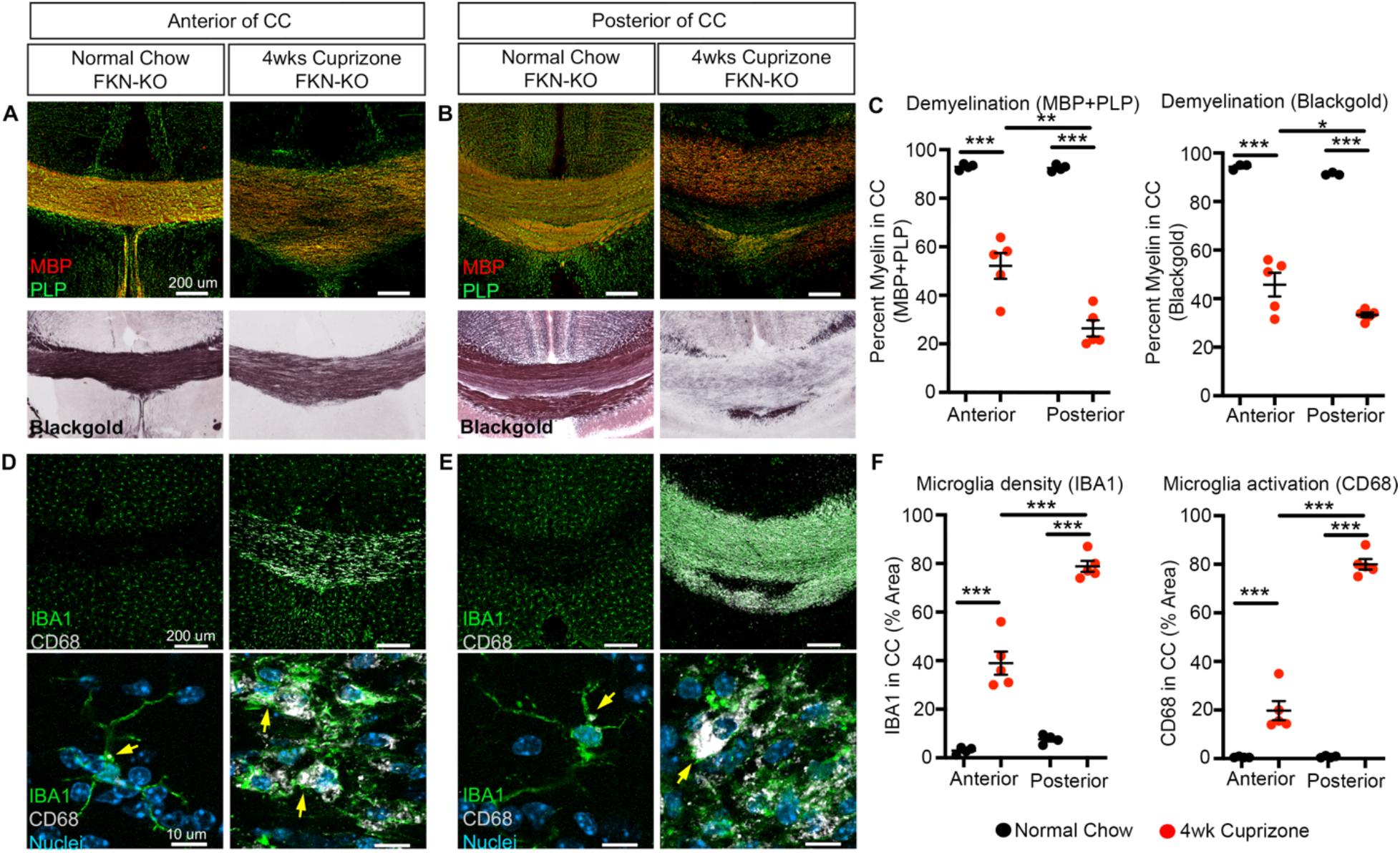
Effects of fractalkine deficiency in cuprizone-induced demyelination. **A-F**, Representative confocal images of brain sections from normal-chow fed and 4 wk cuprizone fed FKN-KO mice, immunostained for (**A, B**) MBP (red) and PLP (green) and blackgold myelin stain or (**D, E**) IBA1 (green) and CD68 (white) in the anterior and posterior of the corpus callosum. **C, F**, Image quantification of normal fed mice (black filled symbols) and 4wk cuprizone fed mice (red filled symbols) is shown. Mean ± SEM, *n* = 4 to 5 mice per group; each dot represents an individual mouse. *P<0.05, **P<0.01, ***<0.001 using one-way ANOVA followed by Tukey’s posttest.

### Proinflammatory gene expression profile is associated with the hCX3CR1^I249/M280^ variant

We next characterized transcriptomic changes in the corpus callosum following acute cuprizone treatment in hCX3CR1^I249/M280^, CX3CR1-KO, and CX3CR1-WT mice using a custom 582 gene inflammatory/immunology panel (Nanostring, see Methods). nCounter digital gene expression analysis identified cuprizone induced 137 differentially expressed genes (DEGs) with 119 DEGs being upregulated relative to normal fed chow mice across all genotypes (Fig. 4A). Interestingly, the cuprizone-induced gene profile was enriched in hCX3CR1^I249/M280^ mice compared to CX3CR1-KO, and CX3CR1-WT mice (Fig. 4A). Consistent with previous reports (Berard et al., 2012; Olah et al., 2012; Clarner et al., 2015), the top five genes upregulated in response to cuprizone were *Itgax, Lcn2, Lilrb4, Cxcl10*, and *Ccl3* and were also highly expressed in tissue extracts from hCX3CR1^I249/M280^ mice (Fig. 4B). As expected, *Mbp* gene expression was significantly downregulated across all three cuprizone-treated genotypes (Fig. 4C). Heat maps depicting normalized gene expression revealed the relative contribution of hCX3CR1^I249/M280^ compared to CX3CR1-KO, and CX3CR1-WT mice on transcriptomic changes in the corpus callosum (Fig. 4D). hCX3CR1^I249/M280^ mice showed increased gene expression for well-known disease-associated microglia markers (*Trem2, Tyrobp, B2m*), prooxidant (*Cybb, Ncf4*), complement system (*Itgax, Itgam, C1qa, C1qb*), Toll-like receptors (*Tlr1, Tlr2, Tlr4, Tlr9, Cd14*) and Fc receptors (*Fcgr4, Fcgr3, Fcgr2b, Fcer1g*) (Gautier et al., 2012; Keren-Shaul et al., 2017; Mendiola et al., 2020) (Fig. 4D). These data suggest that defective fractalkine signaling through hCX3CR1^I249/M280^ triggers microglia proinflammatory and prooxidant transcriptional changes in CNS lesions.

**Figure 4.**
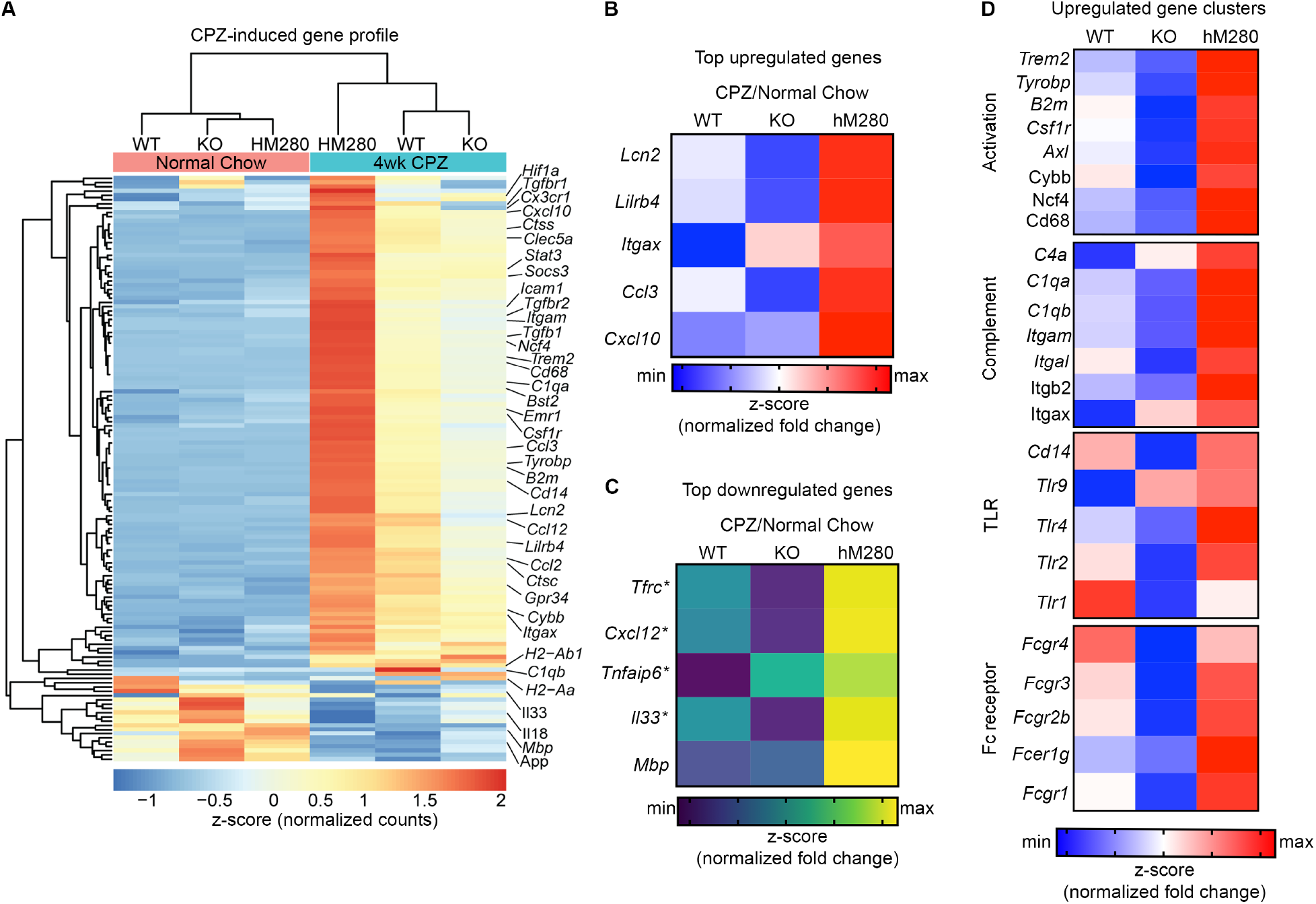
hCX3CR1^I249/M280^ promotes proinflammatory and prooxidant gene expression profiles. **A-D**, Quantitative nCounter gene expression profiling (Nanostring) on corpus callosum tissue extracts from 4wk cuprizone-fed and normal-chow fed hCX3CR1^I249/M280^, CX3CR1-KO and CX3CR1-WT. (A) Heat map of mean normalized gene counts (depicted as z-score expression) of DEGs from cuprizone and normal chow fed mice. Heat maps of top upregulated genes (B), top downregulated genes (C) and upregulated activation gene clusters (D) are shown. Fold changes (respective cuprizone over normal chow) were row normalized and depicted as z-score. Genes with fold change of ≥ 2 and P < 0.05 were deemed as DEG and presented as data mean average of *n* = 3 to 4 mice per group.

### Aberrant CX3CR1-Fractalkine signaling leads to impaired remyelination in hCX3CR1^I249/M280^ mice

We next assessed differences in remyelination by removing cuprizone after four wks and replacing it with normal chow for 1 wk (referred to as 1 wk remyelination). A significant delay in remyelination was observed in the brains of CX3CR1-KO and hCX3CR1^I249/M280^ mice, with evidence of defined demyelinated lesions and decreased MBP/PLP immunoreactivity (Fig. 5A-C, white arrows). While CX3CR1-WT mice recovered up to 75% total myelin in corpus callosum, hCX3CR1^I249/M280^ mice only reached 50% myelin (Fig. 5A-C). Similar results were observed for CX3CR1-KO mice. Furthermore, evaluation of remyelination in the cortex of CX3CR1-KO and hCX3CR1^I249/M280^ mice revealed a reduction in myelin fibers relative to CX3CR1-WT mice (Fig. 5A, bottom panels, white arrows). To highlight impaired remyelination, a side-by-side comparison of the amount of myelin present in the CC at 4wk cuprizone and 1wk remyelination is presented (Fig. 5D). These data suggest aberrant fractalkine-CX3CR1 signaling negatively effects CNS repair mechanisms following a demyelinating event.

**Figure 5.**
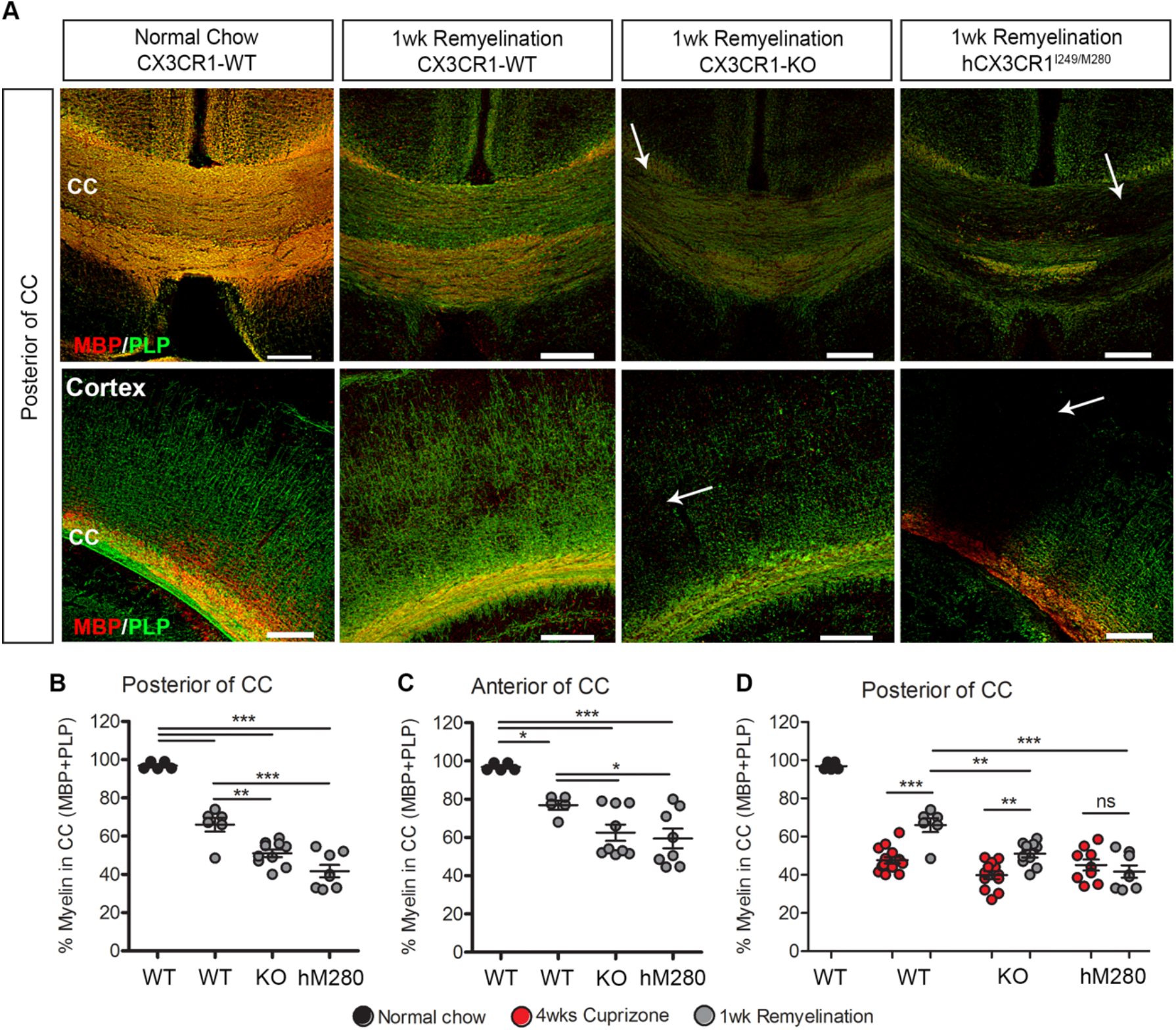
Defective fractalkine signaling delays remyelination in the cuprizone model. **A**, Confocal images of MBP (red) and PLP (green) immunostained brain tissues from 1wk remyelination (4wks cuprizone + 1 wk normal chow) or normal chow fed mice showing the posterior of the corpus callosum and cortex. Image quantification of the extent of remyelination in the (**B**) posterior (**C**) anterior of corpus callosum following 1wk remyelination. (**D**) Side by side comparison of the 4 wk CPZ treated and 1 wk remyelination in the posterior of the CORPUS CALLOSUM is also presented. WT, CX3CR1-WT; KO, CX3CR1-KO; hM280, hCX3CR1^I249/M280^. Data are mean ± SEM, *n* = 5 to 14 mice per group; each dot represents an individual mouse. *P<0.05, **P<0.01, ***P<0.001 by one-way ANOVA followed by Tukey’s posttest. ns = not significant. Scale bars: 200 μm.

### hCX3CR1^I249/M280^ variant dysregulates neurogenesis and oligodendrogenesis

In addition to microglial properties of myelin clearance and remyelination support, microglia are key regulators of neurogenesis and oligodendrogenesis (Shohayeb et al., 2018), however the contribution of hCX3CR1^I249/M280^ remains unknown. To characterize differences in genes regulating neurogenesis in the SGZ in response to focal demyelination in CX3CR1-WT, CX3CR1-KO, and hCX3CR1^I249/M280^ mice, nCounter digital gene expression analysis was performed on SGZ extracts from cuprizone treated mice (Fig. 6). nCounter digital gene expression analysis identified 74 cuprizone induced DEGs (Fig. 6A). The cuprizone-induced gene expression profile for 43 DEGs were upregulated in both CX3CR1-WT and hCX3CR1^I249/M280^ mice compared to CX3CR1-KO mice (Fig. 6A). Consistent with data suggesting that fractalkine signaling may indirectly affect neurogenesis (Fan et al., 2019; Fan et al., 2020), a downregulation in genes associated with neurogenic growth factors (*Nf1, Gdnf, Bdnf, Ptn, Efnb1*), neurogenic enzymes (*Ache, Gpi, Hsp90ab1, Gad65, Sod1*), and transcriptional regulators of neurogenesis (*Neurod1, Hdac4, Kmt2a, Prox1, Ep300, Hey1, Wnt, Tbr2, Pou3f3, Olig2, Apbb1*) was observed in hCX3CR1^I249/M280^ mice (Fig 6C-F).

**Figure 6.**
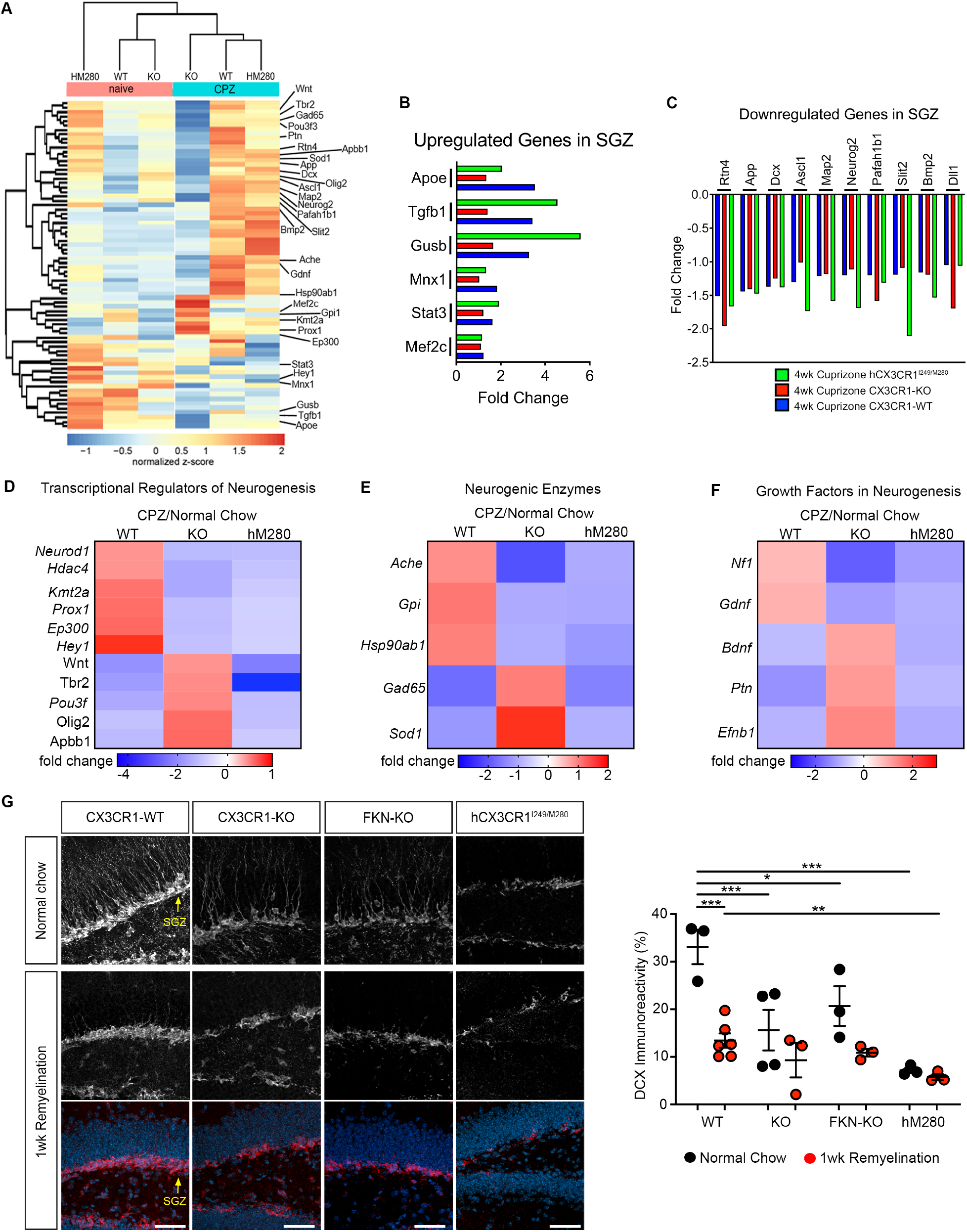
Neurogenesis is impaired in the SGZ of hCX3CR1^I249/M280^ mice. **A-F**, Quantitative nCounter gene expression profiling of neurogenic-related genes (Nanostring) on subgranular zone tissues extracted from 4wk cuprizone-fed and normal-chow fed hCX3CR1^I249/M280^, CX3CR1-KO and CX3CR1-WT mice. (**A**) Heat map of mean normalized gene counts (depicted as z-score expression) of DEGs from cuprizone and normal chow fed mice. Bar graphs depicting the (**B**) top upregulated genes and (**C**) downregulated genes in the SGZ following acute cuprizone treatment are shown. Average fold changes are relative to cuprizone over normal chow genotypes. **D-F**, Heat maps of average fold change expression of (**D**) transcriptional regulators of neurogenesis, (**E**) neurogenic enzymes, and (**F**) growth factors in neurogenesis are shown. Average fold changes are relative to cuprizone over normal chow genotypes. Genes that had a p value < 0.05 were deemed differentially expressed genes (DEGs). Data in **A-F** are geometric means of each transcript for *n* = 3 to 4 mice per group. **G**, Representative images of DCX+ staining (red or pseudocolored white) at the SGZ (yellow arrows) of the hippocampus in coronal brain sections from CX3CR1-WT (WT), CX3CR1-KO (KO), FKN-KO, and hCX3CR1^I249/M280^ (hM280) mice following 1 wk remyelination or in control mice fed normal chow. Nuclei were labeled with Hoechst stain (blue). Percent area of DCX staining was quantified throughout the SGZ for *n* = 3 mice per group and data presented as mean ± SEM. *P<0.05, **P<0.01, ***P<0.001 by one-way ANOVA followed by Tukey’s posttest. Scale bars: 50 μm.

We next validated our gene expression analysis using confocal microscopy and in situ quantification of the abundance of DCX+ neuroblasts throughout the SGZ (Fig. 6G) and mature oligodendrocytes (Fig. 7) in the corpus callosum of experimental animals following remyelination paradigm. In the SGZ, DCX+ neuroblasts and immature neurons exhibited truncated dendrites and were significantly decreased across all groups; however, hCX3CR1^I249/M280^ mice showed the strongest reduction in DCX immunoreactivity after 1 wk remyelination (Fig. 6G). Interestingly, even in normally fed mice a significant reduction in the DCX+ neuroblasts and immature neurons was observed in CX3CR1-KO, FKN-KO and hCX3CR1^I249/M280^ mice (Fig. 6G). Following remyelination, confocal microscopy of IBA1 immunostaining in the corpus callosum showed sustained microgliosis in all genotypes (Fig. 7A). A significant increase in NG2+ cells was across all genotypes compared to untreated mice, with higher counts in hCX3CR1^I249/M280^ mice (Fig. 7B). However, immunohistochemical analyses showed that hCX3CR1^I249/M280^ and CX3CR1-KO mice had significantly fewer CC-1+ mature oligodendrocytes (1301 cells ± 460 and 1629 cells ± 395, respectively) relative to recovering CX3CR1-WT (2731 cells ± 353) and normal-chow fed CX3CR1-WT (3447 cells ± 559) groups (Fig. 7C). Together, these data suggest that fractalkine signaling indirectly influence neurogenesis in general, and is required for efficient OPC differentiation to mature oligodendrocytes during CNS repair.

**Figure 7.**
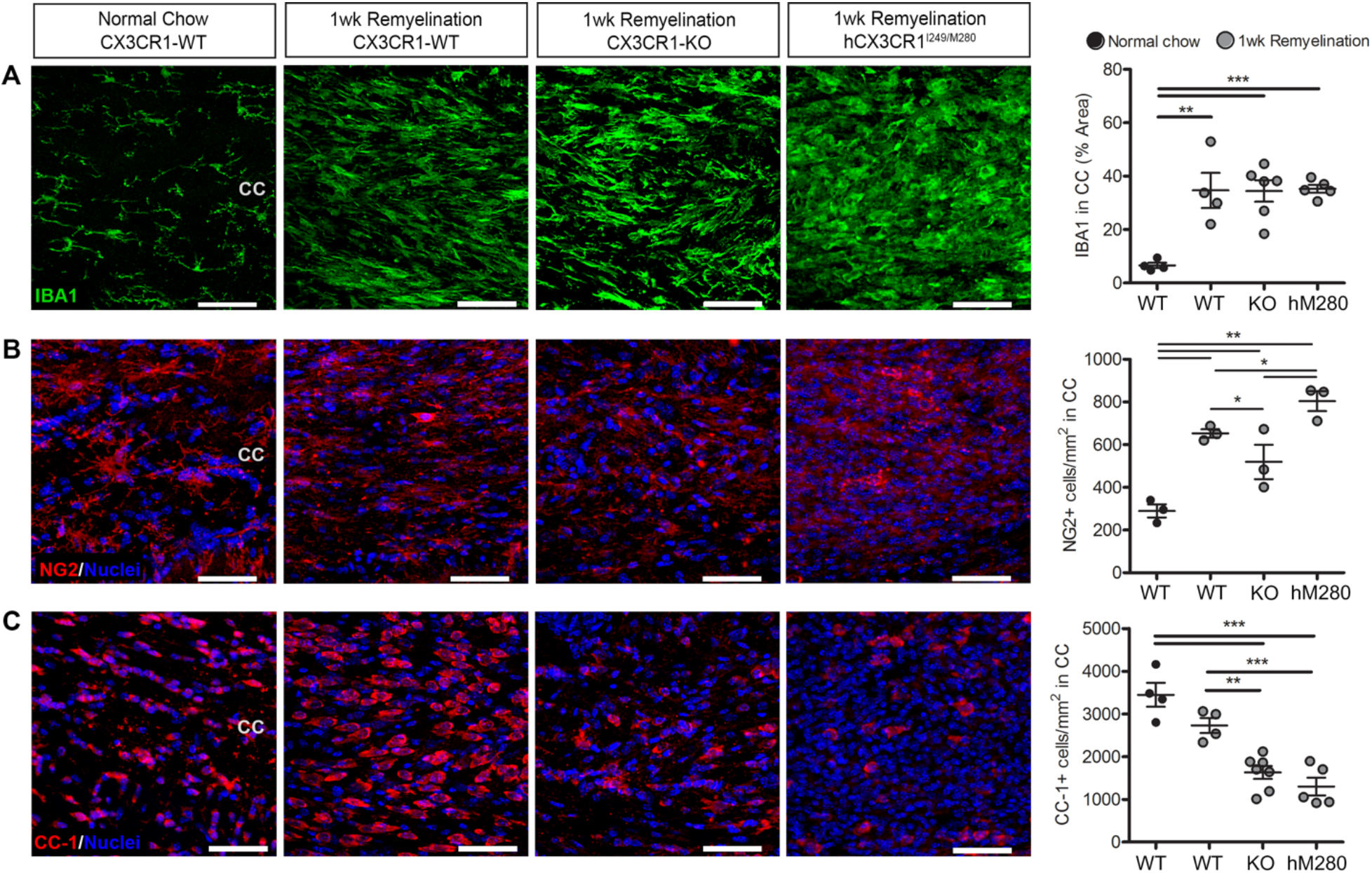
Fractalkine signaling supports oligodendrocyte differentiation following cuprizone-mediated demyelination. **A-C**, Confocal images of (**A**) IBA1+ microglia (green), (**B**) NG2+ cells (red), (**C**) CC-1+ mature oligodendrocytes (red) at the corpus callosum in sections from CX3CR1-WT (WT), CX3CR1-KO (KO), and hCX3CR1^I249/M280^ (hM280) mice following 1 wk remyelination or in control mice fed normal chow. Nuclei were labeled with Hoechst stain (blue). Image quantification is shown on right. Data are mean ± SEM, *n* = 3 to 7 mice per group where each dot represents an individual mouse. *P<0.05, **P<0.01, ***P<0.001 by one-way ANOVA followed by Tukey’s posttest. Scale bars: 50 μm.

## Discussion

This study demonstrates that hCX3CR1^I249/M280^ variant induces aberrant microgliosis in mouse model of demyelination and plays a role in regulating remyelination and generation of progreny from the neurogenic niche. By characterizing mice expressing the hCX3CR1^I249/M280^ variant following cuprizone treatment, we identified that defective CX3CR1 signaling drives inflammatory and oxidative stress gene circuits in CNS lesions, reduces neurogenic precursors and inhibits OPC differentiation. Our results suggest that enhancing CX3CR1 signaling may be important for defining the extracellular milieu by limiting toxic mediators released by microglia that inhibit remyelination and neurogenesis in neurological disorders.

Previous studies have shown that mouse fractalkine is able to signal through hCX3CR1; moreover, receptor mRNA transcript expression in hCX3CR1-M280 brain and spinal cord tissues was comparable to those in WT mice (Cardona et al., 2018). Thus our observations in CPZ model suggest that decreased CX3CR1 signaling contributes to the associated microglial activation phenotype. This work shows that hCX3CR1^I249/M280^ expressing mice, exhibit the most unfavorable remyelination phenotype despite increased NG2+ glia. Lower densities of CC-1+ cells relative to WT mice at 1 wk remyelination, were also observed. Interestingly, similar to naïve and CPZ treated CX3CR1-KO and FKN-KO mice, hCX3CR1^I249/M280^ mice displayed a noticeable decrease in DCX+ cells in the SGZ that mark newly formed and migrating neurons. Distinct from CX3CR1 expressing and deficient mice, the demyelinating phenotype of hCX3CR1^I249/M280^ mice correlated with increased expression of TLR2, shown to restrain remyelination via recognition of hyaluronan oligomers that block OPC maturation and remyelination through TLR2-MyD88 signaling (Sloane et al., 2010); and Lipocalin 2, that with regards to brain autoimmune inflammation was found abundant in CSF of MS patients, and shown to inhibit remyelination in a rat neuroglial cell co-culture model (Al Nimer et al., 2016). Furthermore, phagocytic markers CD68, CD36 and TREM2 were also highly upregulated in hCX3CR1^I249/M280^ tissues suggesting that the remyelination defects were not due to improper housekeeping properties by hCX3CR1^I249/M280^ expressing microglia. Together, these results support the hypothesis that fractalkine/CX3CR1 signaling regulates newly formed neurons and/or the transition of oligodendrocyte progenitors to myelinating oligodendroglia. How or whether new born neurons in the SGZ impact remyelination in the corpus callosum is not clear but has the potential of contributing to cognitive dysfunction.

Cognitive impairments are common clinical manifestations of MS (Rocca et al., 2018). Neuropathological studies have shown demyelination and neuronal loss in the hippocampus of MS patients (Dutta et al., 2011), the brain region with key roles in learning and memory and neurogenesis. A recent study identified hippocampal neurogenesis inversely correlates with the extent of CPZ-induced demyelination (Zhang et al., 2020). Indeed, our data suggest that FKN/CX3CR1 signaling contributes to the proliferation of NSC following demyelination. Our study is in line with another report showing membrane-bound FKN induces neurogenesis in model of Alzheimer’s disease (Fan et al., 2019). Thus, it is possible that neuron-microglial interactions via FKN/CX3CR1 signaling is needed for the restoration of neural circuit formation following a demyelinating event. Future studies are required to identify the precise molecular mechanisms linking focal demyelination with aberrant neurogenesis.

The study presented here is in agreement with published data highlighting remyelination defects in the absence of CX3CR1 signaling (Lampron et al., 2015). However, it has been reported that CX3CR1-KO mice are resistant to cuprizone-induced demyelination in the corpus callosum due to phagocytic defects by microglia and their associated inability to clear myelin debris (Lampron et al., 2015). In order to validate the findings of increased demyelination in CX3CR1-KO mice, we treated FKN-KO mice with cuprizone and observed a similar phenotype in these ligand-deficient mice. Overall in our study, CX3CR1-KO, FKN-KO, and hCX3CR1^I249/M280^ mice were susceptible to cuprizone-induced demyelination without any major differences in the expression of phagocytic markers compared to cuprizone-treated CX3CR1-WT mice.

Although fractalkine/CX3CR1 antagonism is beneficial in some pathologies, the hCX3CR1^I249/M280^ model described in this study will be valuable to address functional differences in CX3CR1 variants. Thus we need to clarify whether the observed differences in the models mentioned above are indeed due to CX3CR1 mediated signaling or to abnormal CX3CR1 function due to desensitization. Moreover, CX3CR1 deficiency delays functional maturation of postsynaptic glutamate receptors in the mouse somatosensory cortex (Hoshiko et al., 2012); leads to impaired hippocampal cognitive function and synaptic plasticity (Rogers et al., 2011); and results in deficits in synaptic pruning, weak synaptic transmission and decreased functional brain connectivity during adulthood (Zhan et al., 2014). Therefore, the signaling via hCX3CR1^I249/M280^ in neuronal processes may provide new avenues to better understand the impact of fractalkine/CX3CR1 antagonist modalities. Nevertheless, these paradoxical observations in CX3CR1-KO models highlight complex functions of CX3CR1 in immune cells versus resident tissue macrophages, and the functional interaction between neurons and microglia via the fractalkine/CX3CR1 axis must be carefully defined before utilization of CX3CR1 antagonistic approaches. Moreover, it is important to clarify the role of fractalkine isoforms, which exist as a membrane-bound and as soluble chemokine, to demyelination demyelination processes in a microglia OPC/OL dependent manner.

The data presented here reveal a decrease in DCX+ immunoreactive area under naive conditions in CX3CR1-KO and hCX3CR1^I249/M280^ mice without significant changes in CPZ-induced demyelination. Although the effect of cuprizone treatment on hippocampal neurogenesis has not been extensively studied, our results agree with previous observations of reduced numbers of DCX+ cells in the cuprizone model (Abe et al., 2015). However, Klein et al found no significant differences in DCX+ cells after 1 or 2 wks remyelination but determined that DCX-expressing neural progenitors can contribute to the mature oligodendrocyte pool (Klein et al., 2020). The observation of increased OPC NG-2+ in hCX3CR1^I249/M280^ mice point to an unexplored area regarding the use of FKN as an effective mediator for the interaction of new neurons and oligodendrocytes to enhance remyelination and repair. Recent studies support the involvement of the fractalkine axis in homeostatic microglia function. Deleting CX3CR1 in the phagocyte system blocks engulfment of OPCs during development which leads to impaired myelin formation (Nemes-Baran et al., 2020). Several critical unanswered questions remain. Detailed analyses of neural lineage and oligodendrocyte lineage progression and distribution in FKN-CX3CR1 expressing, deficient, and hCX3CR1^I249/M280^ mice with a focus in differences in NSC growth factors will be valuable to define the role of the hCX3CR1^I249/M280^ signaling in remyelination and repair. Lastly, identification of differentially regulated genes in microglia that affect cellular number and mobilization of neuroblasts and OPCs in CX3CR1-WT and hCX3CR1^I249/M280^ mice will be valuable to understand further the impact of FKN signaling to subsequent remyelination events.

## Author Contributions

AEC and SMC developed the concept of the study; AEC, ASM designed experiments, analyzed and interpreted data, and wrote the paper. Research development and acquisition of data by ASM, SAM, KC, SMC, DV and IC. AEC, SMC, SVK, DRL and RMR generated the targeting construct for the CX3CR1^I249/M280^ mice. SAL, RMR, SS, EK, CHL and WM interpreted data and assisted in writing the paper. All authors approved the final version of the manuscript.

## Acknowledgements

This study was supported in part by funds from The NIH (SC1GM095426 and R01EY029913 to AEC) and UTSA RISE-PhD Program (Grant GM060655 to ASM and KAC) and by NMSS Postdoctoral Fellowships (FG-1708-28925 to ASM). We acknowledge, the UTSA Cell Analysis Core for supporting this work, and the Genomic & RNA Profiling Core Baylor College of Medicine and their personnel, Lisa D. White, Ph.D, Daniel Kraushaar, Ph.D., Daniela Xavier, Ph.D, Mylinh Bernardi and P30 Digestive Disease Center Support Grant (NIDDK-DK56338) and P30 Cancer Center Support Grant (NCI-CA125123).

## Materials and Methods

### Mice

Male C57BL/6 mice were obtained from Jackson Labs and used as wild type animals (referred to as CX3CR1-WT; RRID: IMSR_JAX:000664). Transgenic CX3CR1-GFP reporter mice (*Cx3cr1*^*GFP/GFP*^; referred to as CX3CR1-KO), hCX3CR1^I249/M280^ (referred to as “hM280” in figures) and *Cx3cl1*^*–/–*^ (referred to as FKN-KO) mice were bred and maintained as previously described (Mendiola et al., 2017; Cardona et al., 2018). This study was carried out in accordance with National Institutes of Health guidelines and approved by The University of Texas at San Antonio Institutional Animal Care and Use Committee.

### Cuprizone-induced demyelination and remyelination model

Seven-to-eight week old male mice were fed 0.2% cuprizone (7012, Red, Rx: 1996698, TD.150233; Envigo) supplemented chow to induce demyelination (Liu et al., 2010). Each animal cage received 50-55 g cuprizone chow and food was replaced every 48 hr until animals were sacrificed. In separate experiments, following cuprizone treatment, food was replaced with normal chow for 1 week and mice were sacrificed to evaluate spontaneous remyelination. In the text, these mice will be referred to as 1wk remyelination. Age and gender matched animals that ate normal chow served as naïve controls.

### Antibodies used in this study

Microglia and macrophages were detected with markers IBA1 (rabbit, 1:4000; Wako, RRID: AB_839504) and CD68 to identify activated phagocytic cells (rat, 1:500; eBioscience, RRID: AB_322219). Myelin detection was done with antibodies against myelin basic protein (MBP; rabbit, 1:4000; Invitrogen, RRID: AB_1501419) and proteolipid protein (PLP; rat, 1:500; Clone A3, obtained from Dr. Wendy Macklin). Mature oligodendrocytes and oligodendrocyte precursor cells (OPCs) were also identified with APC (Ab-7; CC-1; mouse, 1:500, RRID: AB_2057371), and NG2 (Chemicon; rabbit, 1:500, RRID: AB_91789), respectively. Doublecortin-positive neuroblasts and immature neurons were marked with anti-DCX antibody (goat polyclonal, 1:2000; Clone: C-18; Santa Cruz, RRID: AB_2088494). Species-specific secondary antibodies conjugated to Cy3 or Cy5 were purchased from Jackson Laboratories.

### Immunohistochemistry and image quantification

Mice were euthanized under anesthesia by transcardial perfusion with HBSS followed by 4% paraformaldehyde as described previously (Cardona et al., 2015). 30-μm coronal brain sections were generated on a freezing microtome and used for immunohistochemistry. Briefly, 2-3 free-floating sections from the anterior (front: 0.90 – 0.50 mm) and posterior (back: -1.60 – (−2.70 mm) of the corpus callosum of each mouse brain were used. Following antigen retrieval for 15 min at 80**°**C (pH 6.0; DAKO), tissues were incubated in block solution containing 0.3% Triton and 10% goat serum for 1 hr at room temperature (RT) and incubated overnight with primary antibodies at 4**°**C. Fluorescently-conjugated secondary antibodies were used to visualize primary antibodies. Images were acquired on a Zeiss 510 LSM and 3D confocal images were obtained from Imaris software. For image quantification of myelin (MBP and PLP), IBA1 and CD68 signal, raw images were uploaded to ImageJ (NIH), converted to 8-bit grayscale, and then an automatic threshold was applied to the entire image using the plugin otzu thresholding. The intensity of staining was measured as percent area of the corpus callosum. Data were averaged from 2-3 stained sections per region of the corpus callosum and per mouse. DCX+ immunoreactive area was obtained in a similar process with images obtained from the SGZ of the hippocampus. CC-1+ and NG2+ cells were counted in the corpus callosum from 2-3 images per section (2 sections stained per mouse) and normalized to the area. For black-gold histology two free-floating sections from the anterior and posterior of the corpus callosum were stained in a 1.5-mL Eppendorf tube with 500 µL of 0.2% black-gold solution (AG105; EMD Millipore) in a 65**°**C water bath for 12 min. Tissues were then mounted, allowed to dry, and staining was completed according to manufactures instructions. ImageJ analysis was performed similarly as above with thresholding of blackgold staining in each gated corpus callosum kept constant throughout tissues. Data represent percent myelination in the corpus callosum and expressed as black-gold percent area.

### NanoString gene expression analysis

Brains were collected from buffer perfused mice as described above and then the corpus callosum, SGZ and SVZ were separately isolated for RNA extraction. Tissues were homogenized in 1 mL of Trizol reagent (Ambion) and total RNA was extracted according to the manufacturer’s instructions. The integrity and concentrations of RNA were assessed independently (Baylor College of Medicine) using Bioanalyzer (Nanochip, Agilent Technologies). nCounter was used to analyze gene expression in total RNA samples and data were quantified with the nSolver 3.0 software (NanoString Technologies). For corpus callosum gene profiling, the mRNA expression of 582 genes were analyzed using a custom-designed nCounter GE-Mouse Immunology v1 Kit and performed by the Genomic and RNA Profiling Core at Baylor College of Medicine (Department of Advanced Technology Cores). For SGZ and SVZ gene profiling, the mRNA expression of 117 genes were analyzed using a custom-designed nCounter GE-Mouse Neurogenesis Kit and performed by Canopy Biosciences. Using nSolver, all data were initially normalized using positive and negative controls (background subtraction) and housekeeping gene probes. Then treated samples were further normalized to respective normal chow fed genotypes. Differentially expressed genes were defined by a cutoff of ± 2-fold change and P<0.05. Heat maps were generated in R software using ggplot2 and pheatmap packages.

### Quantitative Real-time PCR

RNA was extracted from the corpus callosum as described above and 1 µg of total RNA was reversed transcribed to cDNA using the High-Capacity cDNA Reverse Transcription Kit (Applied Biosystems). Samples were run in triplicate in 10 μL PCR reactions containing 1X SYBR Green PCR Master Mix (Applied Biosystems), 250 nM forward and reverse primers, and 14 ng cDNA template. Results were analyzed by the comparative Ct method, and data are expressed as the average fold change from ΔΔCt for the experimental gene of interest normalized to housekeeping genes (18s and β-actin mRNA) and presented as fold change relative to respective genotype of normally fed mice. The following primers were used: *18s* (Accession number: NR_003278.3): CGGCTACCACATCCAAGGAA, GTCGGAAATACCGCGGTC; *β-actin* (Accession number: NM_007393) CTCTGGCTCCTAGCACCATGAAGA, GTAAAACGCAGCTCAGTAACAGTCCG; *Cd68* (Accession number: NR_110993.1): CTTCCCACAGGCAGCACA, ATTGATGAGAGGCAGCAAGAGG; *Cxcl10* (Accession number: NM_021274.2): TGCTGCCGTCATTTTCTG, GCTCGCAGGGATGATTTCAAG.

### Statistical analysis

Data are expressed as mean ± SEM when scatter plot is provided to show distribution. Otherwise, data are expressed as mean ± SD when described in the Results section and in the Figures. Data were plotted in Graphpad Prism (versions 5 and 9) and statistical tests’ were performed using Student’s t-test when comparing two groups or One-way ANOVA followed by Tukey’s posttest when comparing multiple groups. Statistical significance was deemed when P<0.05.

## References

Abe H, Tanaka T, Kimura M, Mizukami S, Saito F, Imatanaka N, Akahori Y, Yoshida T, Shibutani M (2015) Cuprizone decreases intermediate and late-stage progenitor cells in hippocampal neurogenesis of rats in a framework of 28-day oral dose toxicity study. Toxicol Appl Pharmacol 287:210–221.

Absinta M, Lassmann H, Trapp BD (2020) Mechanisms underlying progression in multiple sclerosis. Curr Opin Neurol 33:277–285.

Al Nimer F, Elliott C, Bergman J, Khademi M, Dring AM, Aeinehband S, Bergenheim T, Romme Christensen J, Sellebjerg F, Svenningsson A, Linington C, Olsson T, Piehl F (2016) Lipocalin-2 is increased in progressive multiple sclerosis and inhibits remyelination. Neurol Neuroimmunol Neuroinflamm 3:e191.

Berard JL, Zarruk JG, Arbour N, Prat A, Yong VW, Jacques FH, Akira S, David S (2012) Lipocalin 2 is a novel immune mediator of experimental autoimmune encephalomyelitis pathogenesis and is modulated in multiple sclerosis. Glia 60:1145–1159.

Cantoni C, Bollman B, Licastro D, Xie M, Mikesell R, Schmidt R, Yuede CM, Galimberti D, Olivecrona G, Klein RS, Cross AH, Otero K, Piccio L (2015) TREM2 regulates microglial cell activation in response to demyelination in vivo. Acta Neuropathol 129:429–447.

Cardona AE, Pioro EP, Sasse ME, Kostenko V, Cardona SM, Dijkstra IM, Huang D, Kidd G, Dombrowski S, Dutta R, Lee J-C, Cook DN, Jung S, Lira SA, Littman DR, Ransohoff RM (2006) Control of microglial neurotoxicity by the fractalkine receptor. Nat Neurosci 9:917–924.

Cardona SM, Mendiola AS, Yang Y-C, Adkins SL, Torres V, Cardona AE (2015) Disruption of Fractalkine Signaling Leads to Microglial Activation and Neuronal Damage in the Diabetic Retina. ASN Neuro 7.

Cardona SM, Kim SV, Church KA, Torres VO, Cleary IA, Mendiola AS, Saville SP, Watowich SS, Parker-Thornburg J, Soto-Ospina A, Araque P, Ransohoff RM, Cardona AE (2018) Role of the Fractalkine Receptor in CNS Autoimmune Inflammation: New Approach Utilizing a Mouse Model Expressing the Human CX3CR1(I249/M280) Variant. Front Cell Neurosci 12:365.

Clarner T, Janssen K, Nellessen L, Stangel M, Skripuletz T, Krauspe B, Hess F-M, Denecke B, Beutner C, Linnartz-Gerlach B, Neumann H, Vallières L, Amor S, Ohl K, Tenbrock K, Beyer C, Kipp M (2015) CXCL10 Triggers Early Microglial Activation in the Cuprizone Model. J Immunol 194:3400–3413.

Cook DN, Chen S-C, Sullivan LM, Manfra DJ, Wiekowski MT, Prosser DM, Vassileva G, Lira SA (2001) Generation and Analysis of Mice Lacking the Chemokine Fractalkine. Mol Cell Biol 21:3159–3165.

Dutta R, Chang A, Doud MK, Kidd GJ, Ribaudo MV, Young EA, Fox RJ, Staugaitis SM, Trapp BD (2011) Demyelination causes synaptic alterations in hippocampi from multiple sclerosis patients. Ann Neurol 69:445–454.

Fan Q, He W, Gayen M, Benoit MR, Luo X, Hu X, Yan R (2020) Activated CX3CL1/Smad2 Signals Prevent Neuronal Loss and Alzheimer’s Tau Pathology-Mediated Cognitive Dysfunction. J Neurosci 40:1133–1144.

Fan Q, Gayen M, Singh N, Gao F, He W, Hu X, Tsai LH, Yan R (2019) The intracellular domain of CX3CL1 regulates adult neurogenesis and Alzheimer’s amyloid pathology. J Exp Med 216:1891–1903.

Garcia JA, Pino PA, Mizutani M, Cardona SM, Charo IF, Ransohoff RM, Forsthuber TG, Cardona AE (2013) Regulation of Adaptive Immunity by the Fractalkine Receptor during Autoimmune Inflammation. J Immunol 191:1063–1072.

Gautier EL, Shay T, Miller J, Greter M, Jakubzick C, Ivanov S, Helft J, Chow A, Elpek KG, Gordonov S, Mazloom AR, Ma’ayan A, Chua WJ, Hansen TH, Turley SJ, Merad M, Randolph GJ, Immunological Genome C (2012) Gene-expression profiles and transcriptional regulatory pathways that underlie the identity and diversity of mouse tissue macrophages. Nat Immunol 13:1118–1128.

Hoshiko M, Arnoux I, Avignone E, Yamamoto N, Audinat E (2012) Deficiency of the Microglial Receptor CX3CR1 Impairs Postnatal Functional Development of Thalamocortical Synapses in the Barrel Cortex. J Neurosci 32:15106.

Keren-Shaul H, Spinrad A, Weiner A, Matcovitch-Natan O, Dvir-Szternfeld R, Ulland TK, David E, Baruch K, Lara-Astaiso D, Toth B, Itzkovitz S, Colonna M, Schwartz M, Amit I (2017) A Unique Microglia Type Associated with Restricting Development of Alzheimer’s Disease. Cell 169:1276–1290 e1217.

Klein B, Mrowetz H, Kreutzer C, Rotheneichner P, Zaunmair P, Lange S, Coras R, Couillard-Despres S, Rivera FJ, Aigner L (2020) DCX(+) neuronal progenitors contribute to new oligodendrocytes during remyelination in the hippocampus. Sci Rep 10:20095.

Kotter MR, Li WW, Zhao C, Franklin RJ (2006) Myelin impairs CNS remyelination by inhibiting oligodendrocyte precursor cell differentiation. J Neurosci 26:328–332.

Lampron A, Larochelle A, Laflamme N, Préfontaine P, Plante M-M, Sánchez MG, Yong VW, Stys PK, Tremblay M-È, Rivest S (2015) Inefficient clearance of myelin debris by microglia impairs remyelinating processes. J Exp Med 212:481–495.

Liu L, Belkadi A, Darnall L, Hu T, Drescher C, Cotleur AC, Padovani-Claudio D, He T, Choi K, Lane TE, Miller RH, Ransohoff RM (2010) CXCR2-positive neutrophils are essential for cuprizone-induced demyelination: relevance to multiple sclerosis. Nat Neurosci 13:319–326.

McDermott DH, Fong AM, Yang Q, Sechler JM, Cupples LA, Merrell MN, Wilson PWF, Agostino RB, Donnell CJ, Patel DD, Murphy PM (2003) Chemokine receptor mutant CX3CR1-M280 has impaired adhesive function and correlates with protection from cardiovascular disease in humans. J Clin Invest 111:1241–1250.

Mendiola AS, Garza R, Cardona SM, Mythen SA, Lira SA, Akassoglou K, Cardona AE (2017) Fractalkine Signaling Attenuates Perivascular Clustering of Microglia and Fibrinogen Leakage during Systemic Inflammation in Mouse Models of Diabetic Retinopathy. Front Cell Neurosci 10.

Mendiola AS et al. (2020) Transcriptional profiling and therapeutic targeting of oxidative stress in neuroinflammation. Nat Immunol 21:513–524.

Miron VE, Boyd A, Zhao JW, Yuen TJ, Ruckh JM, Shadrach JL, van Wijngaarden P, Wagers AJ, Williams A, Franklin RJM, Ffrench-Constant C (2013) M2 microglia and macrophages drive oligodendrocyte differentiation during CNS remyelination. Nat Neurosci 16:1211–1218.

Nemes-Baran AD, White DR, DeSilva TM (2020) Fractalkine-Dependent Microglial Pruning of Viable Oligodendrocyte Progenitor Cells Regulates Myelination. Cell Rep 32:108047.

Olah M, Amor S, Brouwer N, Vinet J, Eggen B, Biber K, Boddeke HWGM (2012) Identification of a microglia phenotype supportive of remyelination. Glia 60:306–321.

Ransohoff RM (2016) How neuroinflammation contributes to neurodegeneration. Science 353:777.

Rocca MA, Barkhof F, De Luca J, Frisen J, Geurts JJG, Hulst HE, Sastre-Garriga J, Filippi M, Group MS (2018) The hippocampus in multiple sclerosis. Lancet Neurol 17:918–926.

Rogers JT, Morganti JM, Bachstetter AD, Hudson CE, Peters MM, Grimmig BA, Weeber EJ, Bickford PC, Gemma C (2011) CX3CR1 Deficiency Leads to Impairment of Hippocampal Cognitive Function and Synaptic Plasticity. J Neurosci 31:16241.

Shohayeb B, Diab M, Ahmed M, Ng DCH (2018) Factors that influence adult neurogenesis as potential therapy. Transl Neurodegener 7:4.

Sloane JA, Batt C, Ma Y, Harris ZM, Trapp B, Vartanian T (2010) Hyaluronan blocks oligodendrocyte progenitor maturation and remyelination through TLR2. Proc Natl Acad Sci U S A 107:11555–11560.

Stojković L, Djurić T, Stanković A, Dinčić E, Stančić O, Veljković N, Alavantić D, Živković M (2012) The association of V249I and T280M fractalkine receptor haplotypes with disease course of multiple sclerosis. J Neuroimmunol 245:87–92.

Voronova A, Yuzwa SA, Wang BS, Zahr S, Syal C, Wang J, Kaplan DR, Miller FD (2017) Migrating Interneurons Secrete Fractalkine to Promote Oligodendrocyte Formation in the Developing Mammalian Brain. Neuron 94:500–516 e509.

Zhan Y, Paolicelli RC, Sforazzini F, Weinhard L, Bolasco G, Pagani F, Vyssotski AL, Bifone A, Gozzi A, Ragozzino D, Gross CT (2014) Deficient neuron-microglia signaling results in impaired functional brain connectivity and social behavior. Nat Neurosci 17:400–406.

Zhang H, Kim Y, Ro EJ, Ho C, Lee D, Trapp BD, Suh H (2020) Hippocampal Neurogenesis and Neural Circuit Formation in a Cuprizone-Induced Multiple Sclerosis Mouse Model. J Neurosci 40:447–458.

